# Reliable detection of translational regulation with Ribo-seq

**DOI:** 10.1101/234344

**Authors:** Sonia P Chothani, Eleonora Adami, Sivakumar Viswanathan, Norbert Hubner, Stuart Cook, Sebastian Schafer, Owen J L Rackham

## Abstract

Ribosome profiling^1^ (Ribo-Seq) reveals genome-wide translation rates via the quantification of ribosome protected fragments (RPFs) of mRNAs. Significant differences of RPFs between conditions can indicate differential translation if they do not match changes in transcription. Reliably identifying translational differences is complicated by the fact that the mRNA level of the transcript directly affects the ribosome occupancy. Changes in mRNA abundances result in changes in RPFs due to differential transcription and not translation. As such, identification of differentially translated genes (DTGs) requires finding significant changes in RPF read counts that cannot be explained by variation in mRNA read counts.

Xiao et al.^2^ developed Xtail with the purpose of more accurately identifying DTGs in pairwise comparisons. Several other methods were also recently released (Ribodiff^3^ and Riborex^4^). At their core, all of these approaches either utilize existing differential expression programs (eg DEseq2^5^ and EdgeR^6^) or use similar statistical assumptions to model the data. However, none of these methods allow for complex experimental design (with more than two conditions) or the use of alternative statistical setups (such as the likelihood ratio tests for comparisons across time) and crucially they do not allow for correction of any batch effects, a commonality to almost all sequencing experiments. Although standalone tools for batch correcting sequencing data exist^7^, differential expression tools for count data require raw read counts to accurately model sample-to-sample variation^8^.

Instead it is possible to tailor the open design of a well established tool, DEseq2 to identify DTGs directly. The introduction of an interaction term provides a coefficient that models non-additive effects of two variables: Condition (untreated/treated) and sequencing methodology (Ribo/RNA). This allows identification of significant differences between conditions that are discordant between sequencing methodologies (see supplementary methods for detailed workflow). This approach can also be extended to facilitate more complex experimental designs, batch effects and other covariates, making DEseq2 a more suitable tool for identifying DTGs. In their paper, Xiao et al. do not include a direct comparison of Xtail and DESeq2; Here, we present a comprehensive benchmarking analysis of all the aforementioned tools to address the same and also evaluate the impact of batch effects on differential translation detection.

We simulated RNA-seq and Ribo-seq data based on published datasets^9^ with a range of batch effects accounting for between 0 to 40% of the variance using Polyester^10^ (figure 1a, supplementary methods). At each batch effect size, simulated Ribo-seq and RNA-seq datasets were generated 20 times and analysed with each of the published methods (our approach, Xtail, Riborex and RiboDiff) to calculate a corresponding set of DTGs. Following this, the associated mean sensitivity and specificity for each method was calculated at various FDR thresholds in order to compare their performance (figure 1b, c, d).

**Figure 1:**
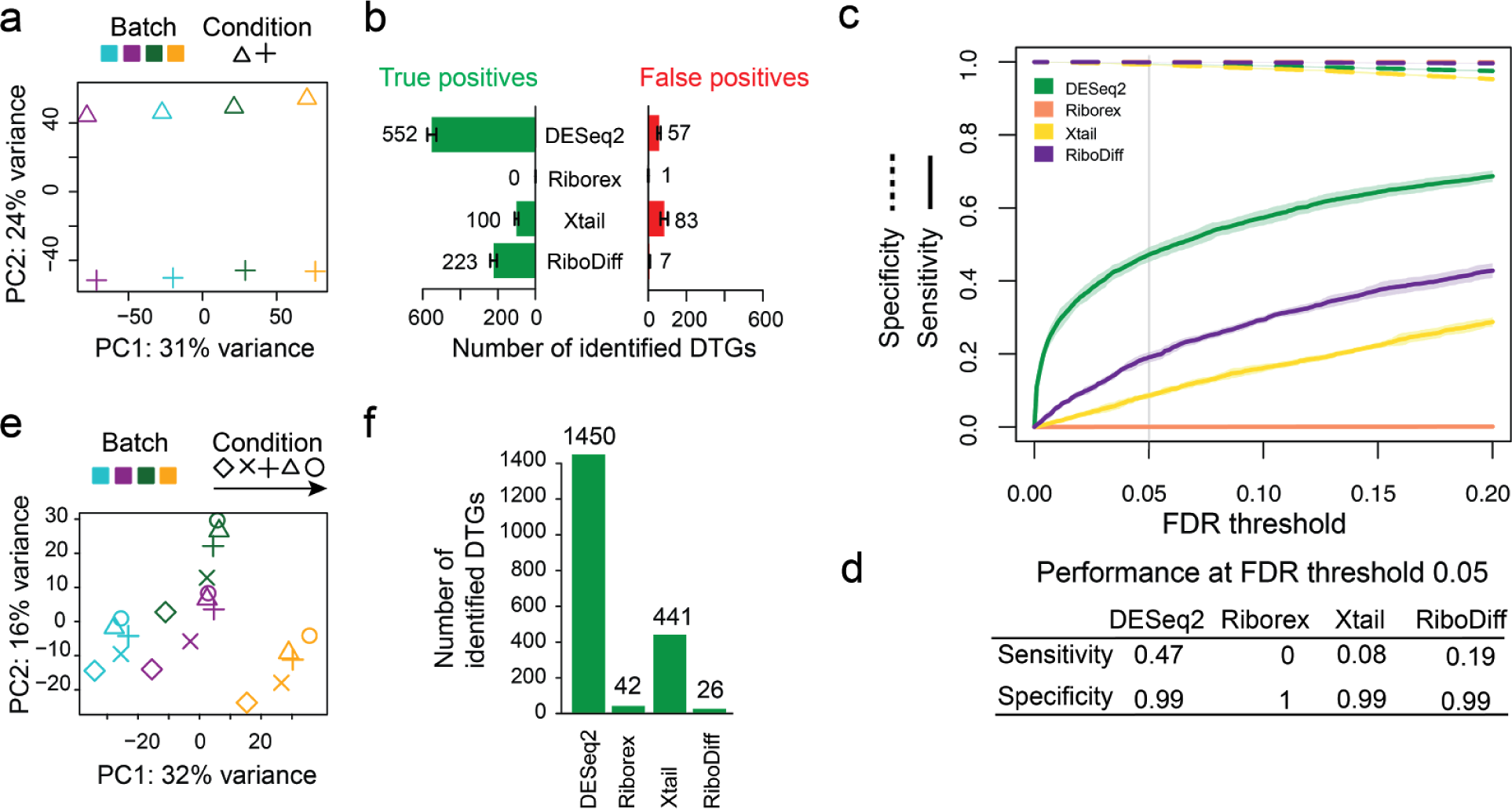
*Detection of differential translation in the presence of batch effects.* *a.-d. Simulated ribosome profiling count dataset with batch effect accounting for 30% of the total variance. a. Principal component analysis. b. Average number of true positive and false positive differentially translated genes (DTGs, to the nearest integer, FDR ≤ 0.05, n=20) found by different methods, c. Resulting sensitivity (True positive rate, solid line) and specificity (True negative rate, dashed line) for different methods over various FDR thresholds. d. Average performance for different methods (FDR ≤ 0.05, n=20). e.-f. Ribosome profiling dataset from primary human fibroblasts in 5 conditions (time series). e. Principal component analysis f. Number of DTGs detected (FDR ≤ 0.05).*

Our tailored approach performs identically or better than all other methods tested in the absence of a batch effect (Supp Fig 1a) and greatly outperforms them in the presence of any batch effect (Supp Fig 1b-h). For instance, in the case of a batch effect that accounts for 30% of the variance and using an FDR cutoff of 0.05, DESeq2 identified 552 true positives with a specificity of >0.99 and Xtail only detects 100 true positives with a similar specificity. Over the range of tested batch effect sizes the sensitivity of DESeq2 drops only by 22.7%, whereas all other methods drop by greater than 80% (Supp Fig 1a-h), making their detection of DTGs substantially less reliable.

To ensure that this effect is not isolated to simulated data, we further analysed a RNA-seq and Ribo-seq dataset from treated and untreated primary human fibroblasts derived from four different donors. In this data we observe a pronounced batch effect between donors, accounting for roughly 30% of the variance in the data (see figure 1d). If this effect is unaccounted for, the sensitivity of Xtail, Riborex and RiboDiff is greatly reduced. The number of significant DTGs across all conditions identified by Xtail (441 genes), Ribodiff (26 genes), Riborex (42 genes) was much lower than our approach that can detect 1450 genes (Figure 1e). It is also important to note here that Xtail takes ~12 hours to execute on this dataset, whilst DESeq2 completes in a matter of seconds.

Since almost all high-throughput sequencing datasets contain batch effects^7^, particularly heterogeneous samples such as human tissues or primary cells, DESeq2-based analysis of differential translation will deliver the most reliable results. Furthermore, recently published methods are inflexible with regards to complex experimental designs such as time-series data and the inclusion of additional covariates, whereas DESeq2 can accommodate all of these situations. We hereby supply the necessary code to perform these analyses with this manuscript.

## Code availability

The code and Rmarkdown files are available on https://github.com/SGDDNB/DTG-detection

## Author contributions

Wet lab experiments were carried out by S.V., E.A. and N.H. Data were analysed and interpreted by S.P.C., S.A.C., S.S. and O.J.L.R. S.P.C., S.S. and O.J.L.R. designed experiments and prepared the manuscript with input from co-authors.

